# Gene cloning and construction of prokaryotic and plant expression vectors of RICIN-A-Chain/PAP-S1 fusion protein and its inhibition of protein synthesis

**DOI:** 10.1101/046060

**Authors:** Yasser S. Hassan, Sherry L. Ogg

**Affiliations:** Research and Development, Ophiuchus LLC, Baltimore, Maryland, U.S.A.; Advanced Academic Programs, Johns Hopkins University, Baltimore, Maryland, U.S.A.

**Keywords:** N-glycosidase, *Phytolacca americana*, pokeweed antiviral protein, Pokeweed antiviral protein from seeds, ribosome-inactivating protein, cDNA, Fusion protein, Inhibition of in vitro protein synthesis

## Abstract

Pokeweed antiviral protein (PAP) is a single-chain ribosome-inactivating protein that exists in several forms isolated from various organs and at different stages of development of *Phytolacca americana* (pokeweed). In this study, PAP-S1, one of the two known isoforms found in seeds, was isolated and PCR amplified using primers based on the known mRNA of PAP-S2, the other known form found in seeds. The complete cDNA encoding PAP-S1 was determined here for the first time. PAP-S1 is a potent antiviral protein with many potential clinical applications. However, it was found to be dosage dependent with observed side effects at high dosage. In this study, we report the production of a recombinant antiviral peptide-fusion protein between Ricin A-chain and PAP-S1. The peptide-fusion recombinant proteins Ricin-A-Chain/PAP-S1 and PAP-S1/Ricin-A-Chain were generated by joining the Nterminus of PAP-S1 to the C-terminus of Ricin A-chain and the C-terminus of PAP-S1 to the N-terminus of Ricin A-chain respectively, and were expressed in an *Escherichia coli* cell free expression systems. The peptide-fusion recombinant protein Ricin-A-Chain/PAP-S1 (F2) was found to be more active than the PAPS1/Ricin-A-chain (F1) and similar to PAP-S1 in a cell free prokaryotic environment, and both showed much stronger activity in a cell free eukaryotic environment. The DNA sequence of the complete cDNA of PAP-S1 and of the peptide-fusion protein Ricin-A-Chain/PAP-S1 with the PAP-S1 signal peptide at the N-terminus of Ricin Achain were inserted in plant destination binary vectors for *A. tumefaciens* mediated transformation. It is the authors’ opinion that additional research should be done in order to determine both cytotoxicity and selectivity of fusion protein F2 compared to PAP-S1, as it could be a viable, more potent and less cytotoxic alternative to PAPS1 alone at high dosage, for both agricultural and therapeutic applications.

## INTRODUCTION

Plants produce toxic proteins known as ribosome inactivating proteins (RIPs) essential to their defense mechanism against outside pathogens. RIPs are RNA N-glycosidases that function by irreversibly inhibiting protein synthesis through the removal of one or more adenine residues from ribosomal RNA (rRNA) [1]. In addition, certain RIPs can remove adenine from DNA and other polynucleotides for which reason they are also known as polynucleotide adenosine glycosidases. Pokeweed antiviral protein (PAP), a RIP from *Phytolacca americana*, can cleave not only adenine, but also guanine from the rRNA of *Escherichia coli*. RIPs are usually categorized into two types, I and II. Type I RIPs are single-chain proteins with a molecular weight of approximately 30 kDa, whereas type II RIPs have an enzymatically active A-chain and a somewhat larger lectin subunit B-chain with a specificity for sugars with galactose-like structures. Type II RIPs have an approximate molecular weight of 56-65 kDa. Given the toxic nature of RIPs, they are usually exported out of the cell once they are synthesized, and are localized either to plant leaves, seeds or roots. Type I RIPs are much more common that type II RIPs and less cytotoxic. This difference in cytotoxicity is believed to be the result of the absence of the cell-binding B chain. However, type I RIPs are still able to enter mammalian cells to some degree in a still poorly understood mechanism. It is hypothesized that they gain access into the cytoplasm as the pathogen enters the cell, thus promoting their activity by impairing host ribosomes. RIPs from plants are being extensively studied as they have potentially useful applications in agriculture and medicine. They have both antiviral and antitumor activities, which have been exploited in the preparation of immunotoxins (via antibody conjugates) by rendering the activities specifically toxic to the targeted cell. The most active areas in biotechnological research into RIPs is targeted at better understanding and subsequent improvement of the cell entry mechanism, increasing specificity, reducing RIP antigenicity, prolonging their plasma half-life and understanding their role in apoptosis [2].

Pokeweed antiviral protein (PAP) is a potent type I RIP expressed in several organs of the plant pokeweed (*Phytolacca americana*), secreted and bound within the plant cell wall matrix [2]. It attacks both prokaryotic and eukaryotic ribosomes and, thus, inhibits protein synthesis. Among the PAP gene family, different genes are expressed in various tissues and at different stages of development in *Phytolacca americana*. PAP, PAP II, PAP-S1, PAP-S2 and PAP-R are the forms that appear in spring leaves, summer leaves, isoform S1 and S2 in seeds, and roots respectively.

The molecular weight ranges from 29 kDa for PAP to 30 kDa for PAP-S’s [3]. PAP-S1 has been identified as the most effective in inhibiting protein synthesis in vitro [4]. PAPs possess antiviral activity on a wide range of plant and human viruses. PAP-S1 and PAP-S2 were found to inhibit infection of HEp-2 cells by herpes simplex virus and poliovirus and also the replication of human immunodeficiency virus 1 (HIV-1) in isolated mononuclear blood cells infected in vitro. It was also found that PAP-S1 and PAP-S2 were more effective than other RIPs at inhibiting the expression of reverse transcriptase in infected cells [3]. It was also found in a recent study that PAP was efficient against Japanese encephalitis virus (same group, family and genus as Zika virus) [5]. It was also observed that expression of the different forms of PAP in transgenic plants leads to broad-spectrum resistance to viral and fungal infections [6][7]. The different forms of PAP have moderate cytotoxicity to non-infected cells and, thus, offer unique opportunities for new applications in therapy and as a protective protein against pathogens in transgenic plants.

Ricin, one of the most potent type II RIPs, produced in the seeds of the castor oil plant, *Ricinus communis*, can efficiently deliver it’s A chain into the cytosol of intoxicated cells through the action of its B chain. The B chain serves as a galactose/N-acetylgalactosamine binding domain (lectin) and is linked to the A chain via disulfide bonds. After the ricin B chain binds complex carbohydrates on the surface of eukaryotic cells containing either terminal N-acetylgalactosamine or beta-1,4-linked galactose residues, it is endocytosed via clathrin-dependent as well as clathrin-independent endocytosis and is thereafter delivered into the early endosomes. It is then transported to the Golgi apparatus by retrograde transport to reach the endoplasmic reticulum (ER) where its disulfide bonds are cleaved by thioredoxin reductases and disulfide isomerases [8]. The median lethal dose (LD50) of ricin is around 22 micrograms per kilogram of body weight if the exposure is from injection or inhalation (1.78 milligram for an average adult). It is important to note that the ricin A chain on its own has less than 0.01% of the toxicity of the native lectin in a cell culture test system. There are no commercially available therapeutic applications of ricin for now, however, ricin A chain is very popular in the development of immunotoxins.

Furthermore, it was shown that ricin A chain alone had no activity on non-infected and tobacco mosaic virus (TMV)-infected tobacco protoplasts alike, while PAPs caused a complete inhibition of TMV production in the infected cells while having no activity on the uninfected protoplasts. Those results led to the conclusion that PAPs, which normally have no mechanism to enter the protoplast, gain entrance to the cytosol of infected cells. It is important to note that in a cell free protein synthesis inhibition assay, it was found that ricin A chain was much more potent than PAPs on rat ribosome inactivation (in the order of 85 fold stronger) [9]. The exact mechanism of the internalization of type I RIPs is still unknown, but this attribute is lacking in the A chain of type II RIPs.

One of the current strategies to develop newer antiviral therapeutics is by creating fusion proteins with higher specificity and selectivity in their interactions leading to less side effects and greater potencies than the proteins on their own. One such example was achieved by Rothan et al. where a peptide-fusion recombinant protein LATA–PAP–THAN (Latarcin-Pokeweed Antiviral Protein I-Thanatin) was found to inhibit Chikungunya virus (CHIKV) replication in the Vero cells at an EC50 of 11.2 lg/ml, which is approximately half of the EC50 of PAP (23.7 lg/ml) and protected the CHIKV-infected mice at the dose of 0.75 mg/ml [10].

It is the objective of this study to create a novel fusion protein between ricin A chain and PAP-S1 in order to achieve greater potency than PAP-S1 alone while keeping its selectivity to infected cells and, in consequence, to reduce previously observed side affects associated to dosage [5]. It was decided to create two versions of the fusion protein, one between ricin A chain C terminus and PAP-S1 N terminus and one between PAP-S1 C terminus and ricin A chain N terminus and determine if they were functional against both prokaryotic and eukaryotic protein synthesis apparatuses.

## MATERIALS AND METHODS

### Materials

The PureLink^®^ Genomic Plant DNA Purification Kit, PureLink^®^ Quick Plasmid Miniprep Kit, HisPur™ Ni-NTA Spin Purification Kit, 0.2 m, Phire Plant Direct PCR Master Mix, PureLink^™^ Quick Gel Extraction and PCR Purification Combo Kit, Phire Hot Start II DNA Polymerase, Gateway^®^ BP & LR clonase II enzyme mix, Gateway^®^ pDonr-221 vector, Gateway^®^ pExpl-Dest vector, Expressway^™^ Mini Cell-Free Expression System, One Shot^®^ TOPIO Electrocomp^™^ *E. coli*, and ElectroMAX^™^ *A. tumefaciens* LBA4404 Cells were purchased from Thermo Fisher Scientific. The pH7WG2D [12] plant destination binary vector for *A. tumefaciens* mediated transformation was purchased from the University of Gent. The oligonucleotides used for PCR, creating attb PCR products and overlap extension PCR products, were synthesized by Integrated DNA Technologies. The Rabbit Reticulate Lysate TnT^®^ Quick Coupled Transcription/Translation System and the *E. coli* S30 T7 High-Yield Protein Expression System were purchased from Promega.

### Amplification of DNA by PCR

#### PAP-S1

Genomic DNA was prepared from seeds of *Phytolacca americana* using the PureLink^®^ Genomic Plant DNA Purification Kit. Since only a partial cds of PAP-S1 (GenBank: AB071854.1) was available, different primers were designed based on the available mRNA PAP-S2 cds (GenBank: X98079.1) to sequence the complete cds of PAP-S1. The assumption that primers based on the mRNA of PAP-S2 would allow us to amplify PAP-S1 sequence was based on the fact that PAP-S1 and PAP-S2 are highly identical, and also on the previous speculation that PAP-S1 and PAP-S2 were simply two different forms of only one available pokeweed antiviral protein present in seeds, and thus in the same locus leading to similar mRNA [3]. The forward primer used was 5’-AGAAGCGGCAAAGGGAAGATGAA-3’ and for the reverse primer was 5’-CCATTGGCCCGGCCTCTTATTT-3’. The sequence of PAP-S1 was amplified by PCR using the Phire Hot Start II DNA Polymerase kit based on the 50ul protocol with the melting temperature of the primers determined by the TM calculator provided by Thermo Fisher Scientific. The PCR products were separated by agarose gel electrophoresis. A fragment with the expected size (about 900 bp) was extracted and used as a template for the second PCR under the same conditions. The products of the second PCR (about 900bp) was purified by agarose gel electrophoresis using the PureLink^™^ Quick Gel Extraction and PCR Purification Combo Kit.

#### Ricin A Chain

The sequence of the ricin A chain was isolated and amplified directly from seeds of *Ricinus communis* using the Phire Plant Direct PCR Master Mix. The complete sequence of ricin is widely available and the primers were designed based on the available *Ricinus communis* ricin gene (GenBank: X52908.1). The forward and reverse primers were selected based on the Integrated DNA Technologies primers design software to amplify ricin A chain including the linker peptide region. The forward primer selected was 5’-GAGAATGCTAATGTATTTGGACAGCCA-3’ and the reverse one 5’-GTATTGCGTTTCCGTTGTGGAATCT-3’. The sequence of ricin A chain with the linker peptide region was purified by agarose gel electrophoresis the same way as described for PAP-S1 (fragment of about 1kb).

## Design of fusion DNA sequence with partial attb sequence

### PCR extension of PAP-S1 for fusion at the N terminus

The N terminus of PAP-S1 was extended by PCR using the following forward primer 5’-CAGTGGTACCAAATTTTAATATAAATACAATCACCTTCGA-3’ (overlapping the Ricin A-chain linker peptide sequence) and reverse primer 5’-AGAAAGCTGGGTAGAACATGGCATTTTGTTA-3’ using the Phire Hot Start II DNA Polymerase kit based on the 50ul protocol. The PCR product was purified by agarose gel electrophoresis as described previously (band at 800bp).

### PCR extension of PAP-S1 for fusion at the C terminus

The C terminus of PAP-S1 was extended using the following forward primer 5’-AAAAAGCAGGCTCTATAAATACAATCACCTTCGA-3’ (without the signal sequence of PAP-S1) and reverse primer 5’-GTACCACTGGCCTTATAAGCAAAGAAGTTGCTTGGCAAGTC-3’ (using the Ricin linker peptide as linker between PAP-S1 C Terminus and Ricin A Chain N Terminus) using the Phire Hot Start II DNA Polymerase kit based on the 50ul protocol. The PCR product was purified by agarose gel electrophoresis as described previously (band at 800bp).

### PCR extension of Ricin A-chain at the C terminus

The C terminus of Ricin A-chain was extended by PCR with the following forward primer 5’-AAAAAGCAGGCTCTATATTCCCCAAACAATAC-3’ and reverse primer 5’-TCGAAGGTGATTGTATTTATATTAAAATTTGGTACCACTG-3’ (overlapping PAP-S1 protein cds) using Phire Hot Start II DNA Polymerase kit based on the 50ul protocol. The PCR product was purified by agarose gel electrophoresis (band at 850bp).

### PCR extension of Ricin A-chain at the N terminus

The N terminus of Ricin A-chain was extended by PCR with the following forward primer 5’-GCTTATAAGGCCAGTGGTACCAAATTTTAATATATTCCCCAAACAATAC −3’ (overlapping the ricin linker peptide) and reverse primer 5’-GAAAGCTGGGTTAAAACTGTGACGATGGT-3’ using Phire Hot Start II DNA Polymerase kit based on the 50ul protocol. The PCR product was purified by agarose gel electrophoresis (band at 850bp).

### Fusion protein sequence

#### At the C terminus of Ricin

The C terminus of Ricin A-chain (including the linker peptide region) was fused to the N terminus of PAP-S1 (without the signal peptide) by overlap extension PCR.

The forward and reverse primers were 5’-AAAAAGCAGGCTCTATATTCCCCAAACAATAC-3’ and 5’-AGAAAGCTGGGTAGAACATGGCATTTTGTTA-3’ respectively using Phire Hot Start II DNA Polymerase kit based on the 50ul protocol. The PCR product was purified by agarose gel electrophoresis (band at 1650bp).

### At the C terminus of PAP-S1

The N terminus of Ricin A-chain (with the linker peptide region fused at the N terminus) was fused to the C terminus of PAP-S1 (by overlapping the linker peptide region) by overlap extension PCR. The forward and reverse primers were 5’-AAAAAGCAGGCTCTATAAATACAATCACCTTCGA-3’ and 5’-GAAAGCTGGGTTAAAACTGTGACGATGGT-3’ respectively using Phire Hot Start II DNA Polymerase kit based on the 50ul protocol. The PCR product was purified by agarose gel electrophoresis (band at 1650bp).

## DNA Sequencing

The nucleotide sequences of inserts cloned into the two different plasmids DNA were determined using multiple primers using Sanger DNA Sequencing by an outside laboratory facility (GENEWIZ).

### Production of Recombinant PAP-S1 and Fusion proteins

#### Construction of prokaryotic expression vectors

The PAP-S1 and Fusion Protein DNA sequence were each inserted into the pexpl-Dest vector using the Gateway System One Tube format from Thermo Fisher Scientific (combining the BP and LR reactions). The PAP-S1 and Fusion Protein DNA sequences were flanked with the attBl and attB2 sequences (by adding them to the PCR forward and reverse primers, respectively) prior to the One Tube format. This step is essential as the BP and LR recombination reactions allow the attB flanked PCR product to be inserted into an attP containing donor vector to generate an attL flanked insert entry clone (proprietary enzyme mix BP Reaction), and this attL flanked insert in the entry vector is then inserted into the attR destination vector (proprietary enzyme mix LR reaction).

#### Construction of plant expression vectors

The PAP-S1 and Fusion Protein (at the N terminus of PAP-S1) DNA sequences were each inserted into the pH7WG2D plant destination binary vector using the Gateway System One Tube format from Thermo Fisher Scientific (combining the BP and LR reactions). The signal peptide sequence of PAP-S1 was first added at the N terminus of the PAP-S1 and Fusion Protein DNA sequences by extension overlap PCR as previously described. The PAP-S1 and Fusion protein DNA sequence were then flanked with the attB1 and attB2 sequences (by adding them to the PCR forward and reverse primers, respectively) prior to the One Tube format.

#### Cloning of expression plasmids

Both PAP-S1 and Fusion prokaryotic and plant expression plasmids were transfected by electroporation using an Eppendorf Eporator^®^ into One Shot^®^ T0P10 Electrocomp^™^ *E. coli* for plasmid propagation. The plasmids were then purified using the PureLink^®^ Quick Plasmid Miniprep Kit. The plasmids were ethanol precipitated and concentrated to 500ng/ul.

#### Prokaryotic expression

Prokaryotic expression of both PAP-S1 and Fusion proteins were achieved using the Expressway^™^ Mini Cell-Free Expression System. The proteins were purified using the HisPur^™^ Ni-NTA Spin Purification Kit, 0.2 m before being run on protein gels for confirmation.

#### Inhibition of cell-free protein synthesis

The enzyme activity of the purified recombinant proteins was determined by intensity of the band on protein gel of a control against expression of the control without the recombinant proteins, after protein purification using the HisPur^™^ Ni-NTA Spin Purification Kit, 0.2 m, in both the Rabbit Reticulate Lysate TnT^®^ Quick Coupled Transcription/Translation System and the *E. coli* S30 T7 High-Yield Protein Expression System. The control used was pEXP5-NT/CALML3 control vector as DNA template expressing an N-terminally-tagged human calmodulin-like 3 (CALML3) protein (under the T7 promoter). The concentration of CALML3 was determined for increasing concentrations of recombinant PAP-S1 and Fusion proteins by visually measuring band intensity on a protein gel after Coomassie blue staining.

## RESULTS AND DISCUSSION

### PAP-S1

It was difficult to extract useful genomic DNA from seeds of *P. americana* given the small size of its seeds and its fibrous nature. Additionally, the reverse primer led to many unspecific binding events, and gave unwanted PCR products. The complete cds of PAP-S1 was however determined and is now available at GenBank with the accession number KT630652. The signal peptide DNA sequence of PAP-S1 is 83% identical to the signal peptide of PAP-S2, however, the translated proteins are only 67% identical based on Blast (see below Figure 1 for both the nucleotide and protein alignment between the determined PAP-S1 and the PAP-S2 signal peptide as reported by Poyet et al.). These differences in amino acids might be significant since the signal peptide has been shown to play roles in toxicity, specificity and expression of RIPs [13] and, thus, might shed new light on those particular mechanisms.

**Figure 1.**
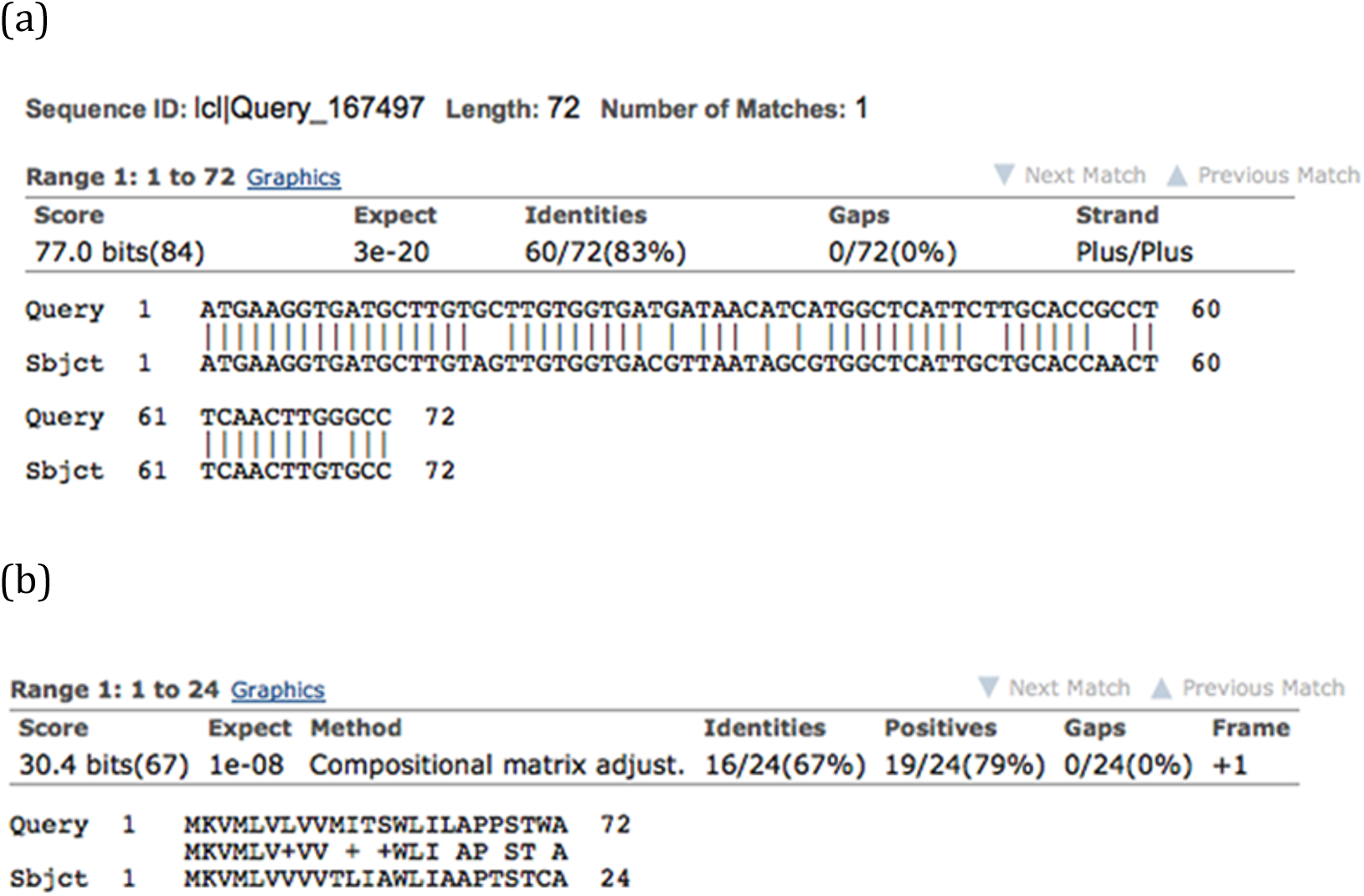
Alignment of the two signal peptide sequences using blastn and blastp. (a) Blastn of the two signal peptide sequences have an identities score of 83%, however, (b) Blastp of the translated nucleotide sequences have only an identities score of 67%.

### Expression Vectors and Production of recombinant Proteins in Cell Free Expression System

The prokaryotic and eukaryotic expression vectors were constructed following the Gateway cloning system, and no major difficulties were encountered. The DNA structures of the expression vectors used for prokaryotic expression are shown in Figure 2 and in Figure 3 for plant expression using binary vectors.

**Figure 2.**
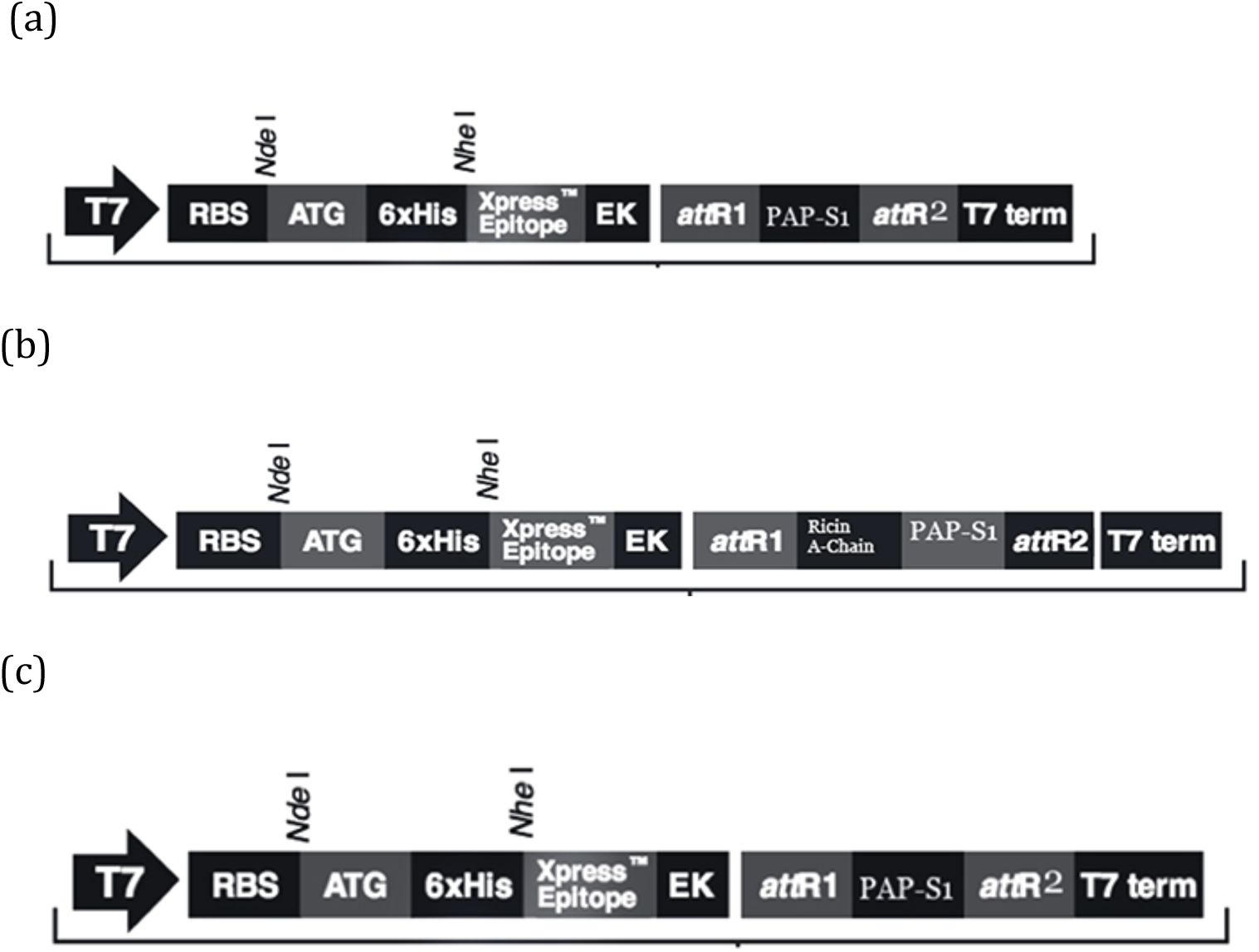
DNA structures for the expression vectors of (a) PAP-S1, (b) Ricin-A-chain/PAP-S1 fusion protein and (c) PAP-S1/Ricin A-chain.

**Figure 3.**
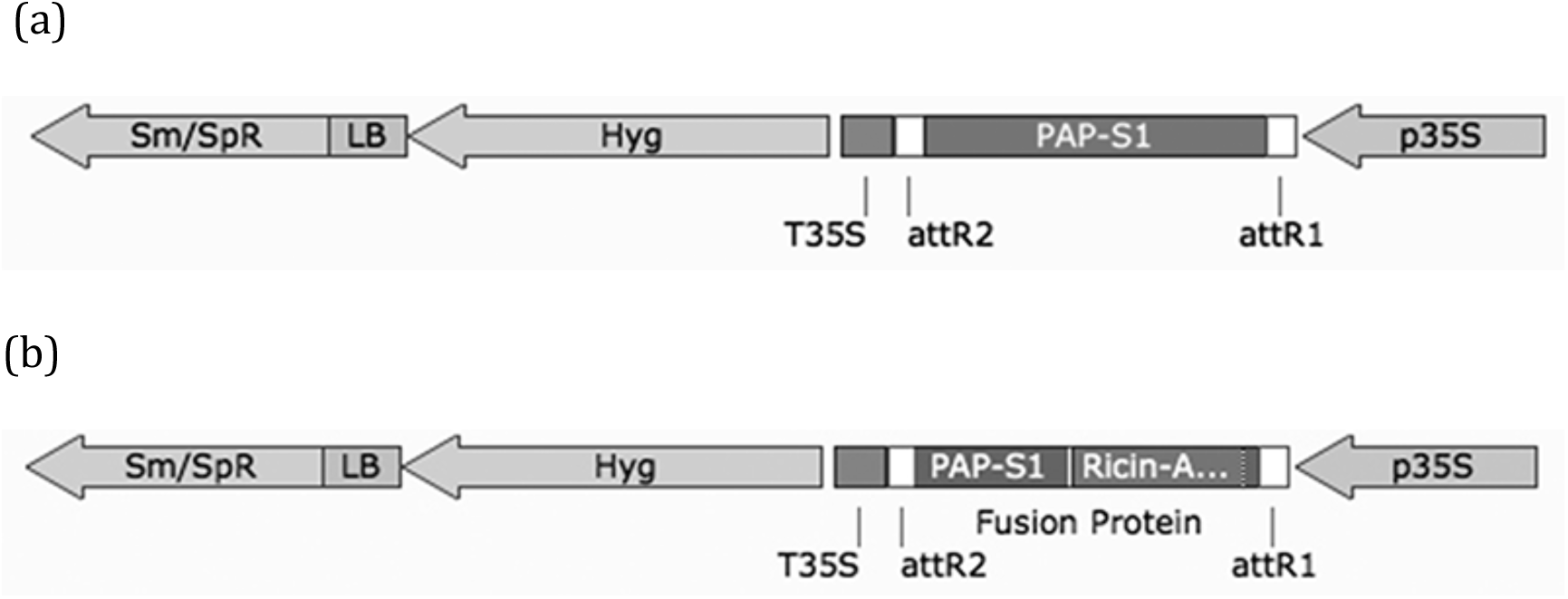
T-DNA structure of the binary vector pH7WG2D constructed for plant expression mediated by *A. tumefaciens* transformation.

Arrows indicate transcription orientation. T7 is the T7 promoter, RBS is the Ribosome binding site, ATG is the ATG start codon, 6xHis is the Polyhistidine region, Xpress^™^ epitope is as stated, EK is the Enterokinase recognition sit, attR1 is the resulting attR1 site from the LR clonase reaction, (a) PAP-S1 is the PAP-S1 insert without its signal peptide, attR2 is the attR2 site resulting from the LR clonase reaction, T7 is the T7 transcription termination region. (b) Ricin A-chain is the Ricin A-chain insert without its signal peptide but with the linker peptide region at its C terminus, PAP-S1 is PAP-S1 insert with the ricin-A-chain linker peptide region at its N terminus.(C) Ricin A-chain is the Ricin A-chain insert without its signal peptide but with its linker peptide region at its N terminus, PAP-S1 is the PAP-S1 insert with the ricin-A-chain linker peptide region at its C terminus.

Arrows indicate transcription orientation. LB is the left border; P35S and T35S, CaMV35S promoter and terminator; Hyg is the Hygromycin resistance gene, Sm/SpR is the Spectinomycin resistance gene, (a) PAP-S1 is the PAP-S1 insert with its signal peptide, (b) PAP-S1 Ricin-A … is the Ricin-A-chain/PAP-Sl insert with the PAP-S1 signal peptide at the N terminus of Ricin-A-Chain and the linker peptide region of Ricin A-chain at its C terminus and N terminus of PAP-S1.

However, it was observed that it was very difficult to obtain high concentration of usable plasmids in sufficient quantities even after an overnight culture for prokaryotic expression due to the high instability of the proteins expressed. Indeed, even when the recombinant proteins were expressed in minimal amounts due to the leaking nature of the vector, it was found that they were extremely toxic to E. *coli* as was previously observed [14]. For this reason, PCR amplifications of the genes of interests from the T7 promoter to the T7 terminator regions were done from the plasmids. Linear DNA (including the control) was used in the Expressway Mini Cell free System to generate the recombinant proteins. The proteins were expressed and visible bands showed up where expected on protein gels stained with Coomassie blue (Figure 4), but it is important to note that the levels of expression for the three recombinant proteins were very low compared to the control protein and, more importantly, the level of expression of the fusion protein at the PAP-S1 C terminus (F1) was higher than that of the fusion protein at PAP-S1 N terminus (F2). The expression levels of PAP-S1 were the lowest and barely visible. Recombinant proteins PAP-S1 and F1 were thus expressed in thrice the reaction volume (150ul). More visible bands can be seen in Figure 5. The low levels of expression of the recombinant proteins were anticipated since the recombinant proteins were expected to be toxic to prokaryotic ribosomes. PAP-S1 is highly toxic to prokaryotic ribosomes as it can cleave not only adenine, but also guanine from the rRNA of *Escherichia coli* as above mentioned. In regard to F1 and F2, the recombinant proteins were successfully produced and their low expression levels show that they are both functional; F2 being more potent than F1 and PAP-S1 more potent than F2 in a prokaryotic environment. This was expected since Ricin A chain is known to be inactive against prokaryotic ribosomes.

**Figure 4.**
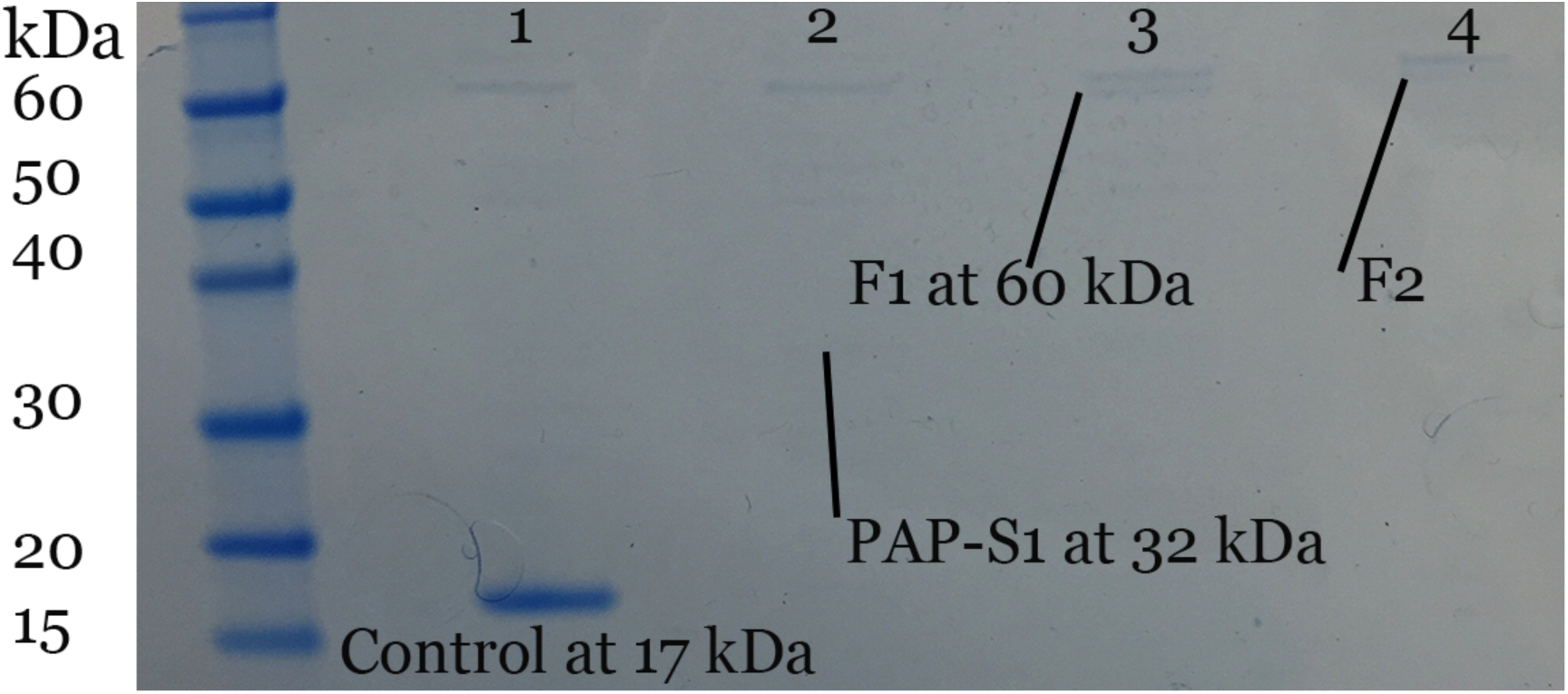
Recombinant proteins gel stained with Coomassie blue after His-Tag purification. Lane 1: Control protein after His-Tag Purification. The band is clearly visible at 17kDa. Lane 2: PAP-S1 recombinant protein barely visible at the 32kDa line. Lane 3: F1 (at the PAP-S1 C terminus) recombinant protein clearly visible at the 60kDa line. Lane 4: F2 (at the PAP-S1 N terminus) recombinant protein visible at the 60kDa line, but less pronounced than the F1 band. ^*^All other bands are due to proteins going through the His-Tag purification column from the initial expression reaction.

**Figure 5.**
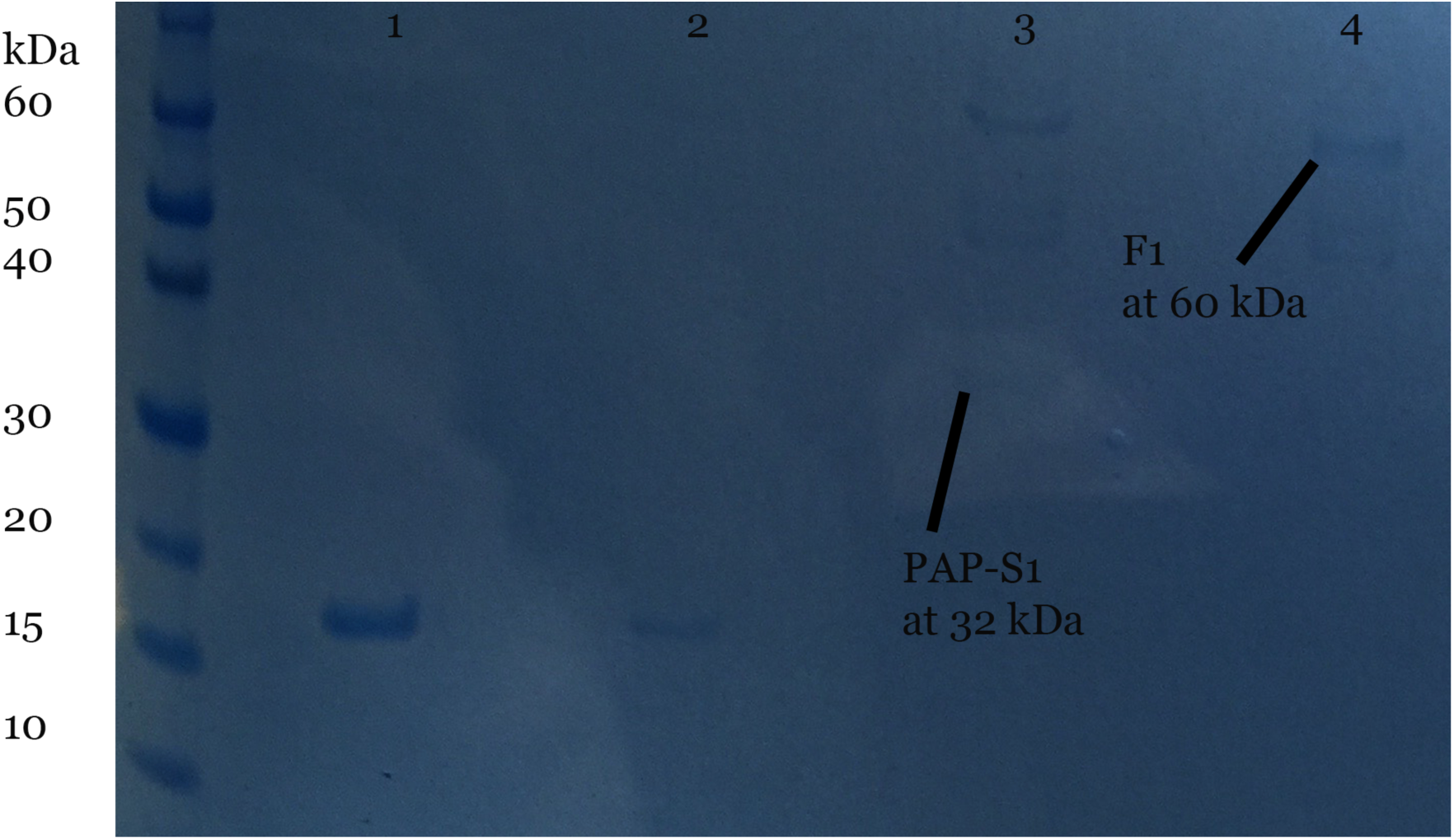
Gel stained with Coomassie blue after His-Tag purification of recombinant protein PAP-S1 in 3X volume reaction. Lane 1: Control protein (expressed from circular DNA) after His-Tag Purification. The band is visible at 17kDa. Lane 2: Control protein (expressed from linear DNA) after His-Tag Purification. The band is less visible at 17 kDa than when expressed from circular DNA as predicted by manufacturer. Lane 3: PAP-S1 recombinant protein visible at 32 kDa line. Lane 4: F1 recombinant (at the PAP-S1 C terminus) protein visible at 60 kDa line. ^*^All other bands are due to proteins going through the His-Tag purification column from the initial expression reaction.

### Activity of Recombinant F1, F2 and PAP-S1

In order to have a better idea of the activity of the recombinant proteins, protein inhibition synthesis assays in both prokaryotic and eukaryotic systems were achieved using various kits. The same volume of PAP-S1, F1 and F2 were then used in two different prokaryotic systems, the *E. coli* S30 T7 High-Yield Protein Expression System and the Expressway^™^ Mini Cell-Free Expression System. It is important to note that the expression of F1 was shown to be greater than both F2 and PAP-S1 in the cell free expression system. Thus, 20% volume of F1 would be more concentrated than 20% F2 and than 20% PAP-S1. Similar results were observed where low volume of PAP-S1, F1 and F2 showed some protein inhibition (Lanes 3, 4 and 5 from Figure 6 and 7 below). In order to determine which of F1 or F2 is the most active, increasing volumes of F1 and F2 were used against control DNA. It was observed that F2 is more active than F1 with increasing concentrations, and even comparable to PAP-S1. Figure 8 shows 25% volume of F2 to be as potent as 36% volume of F1, and 36% volume of F2 almost completely inhibits protein synthesis while F1 at 36% still shows a visible band at 17kDa. This difference in activity might be due to the C terminus of PAP-S1 being free in F2 as it was previously observed that the C terminus has distinct roles in transport to the cytosol, ribosome depurination and cytotoxicity [11]. It could also be due to a conformation difference between F1 and F2.

**Figure 6.**
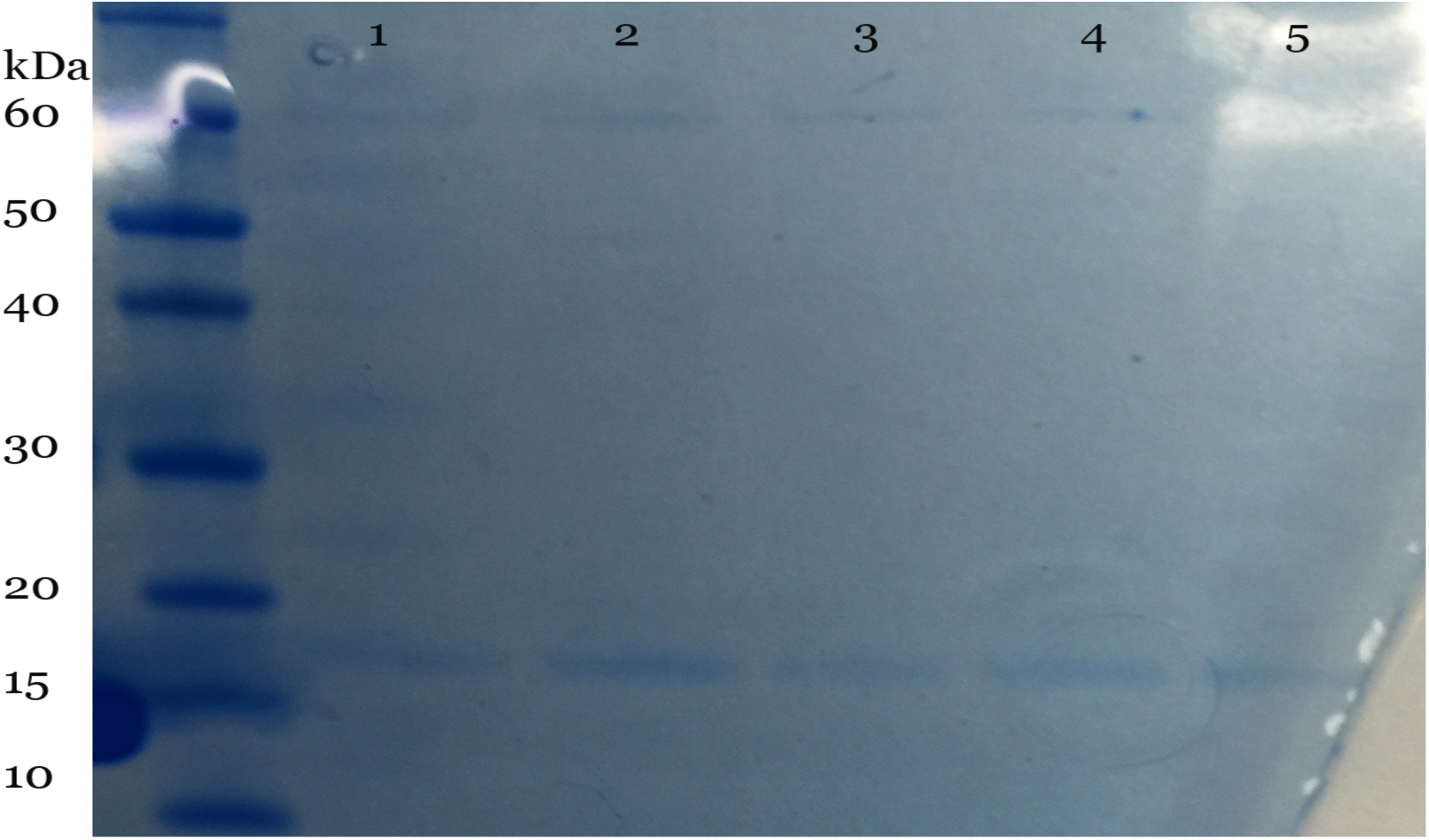
Inhibition of protein synthesis by recombinant proteins in *E. coli* S30 T7 High-Yield Protein Expression System Lane 1: Control DNA at 17 kDa with >250 mM Imidazole to verify impact on protein synthesis Lane 2: Control DNA at 17kDa Lane 3: PAP-S1 20% volume of reaction (50ul) (10ul of His-Tag purified PAP-S1) Lane 4: F1 20% volume (10ul of His-Tag purified F1) Lane 5: F2 20% volume (10ul of His-Tag purified F2) ^*^All other bands are due to proteins going through the His-Tag purification column from the initial expression reaction.

**Figure 7.**
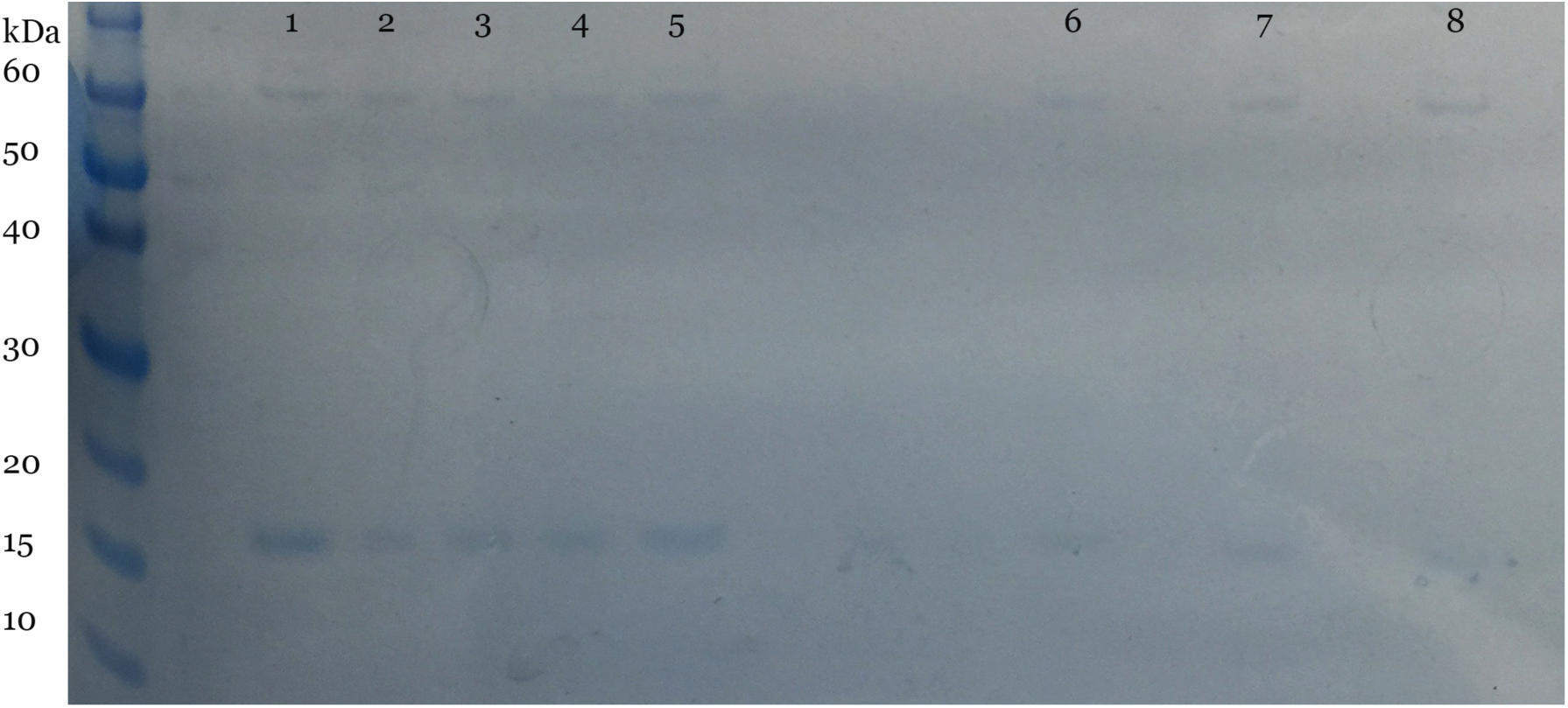
Inhibition of protein synthesis by recombinant proteins in the Expressway^™^ Mini Cell-Free Expression System. Lane 1: Control DNA at 17kDa Lane 2: Control DNA at 17 kDa with >500 mM Imidazole to verify impact on protein synthesis Lane 3: PAP-S1 11.5% volume of reaction (50ul reaction + 50ul buffer) (5.75ul of His-Tag purified PAP-S1) Lane 4: F1 11.5% volume (5.75 ul of His-Tag purified F1) Lane 5: F2 11. 5% volume (5.75ul of His-Tag purified F2) Lane 6: F2 25% volume (15ul of His-Tag purified F2) Lane 7: F1 36% volume (25ul of His-Tag purified F1) Lane 8: F2 36% volume (25ul of His-Tag purified F2) ^*^All other bands are due to proteins going through the His-Tag purification column from the initial expression reaction or spillovers.

**Figure 8.**
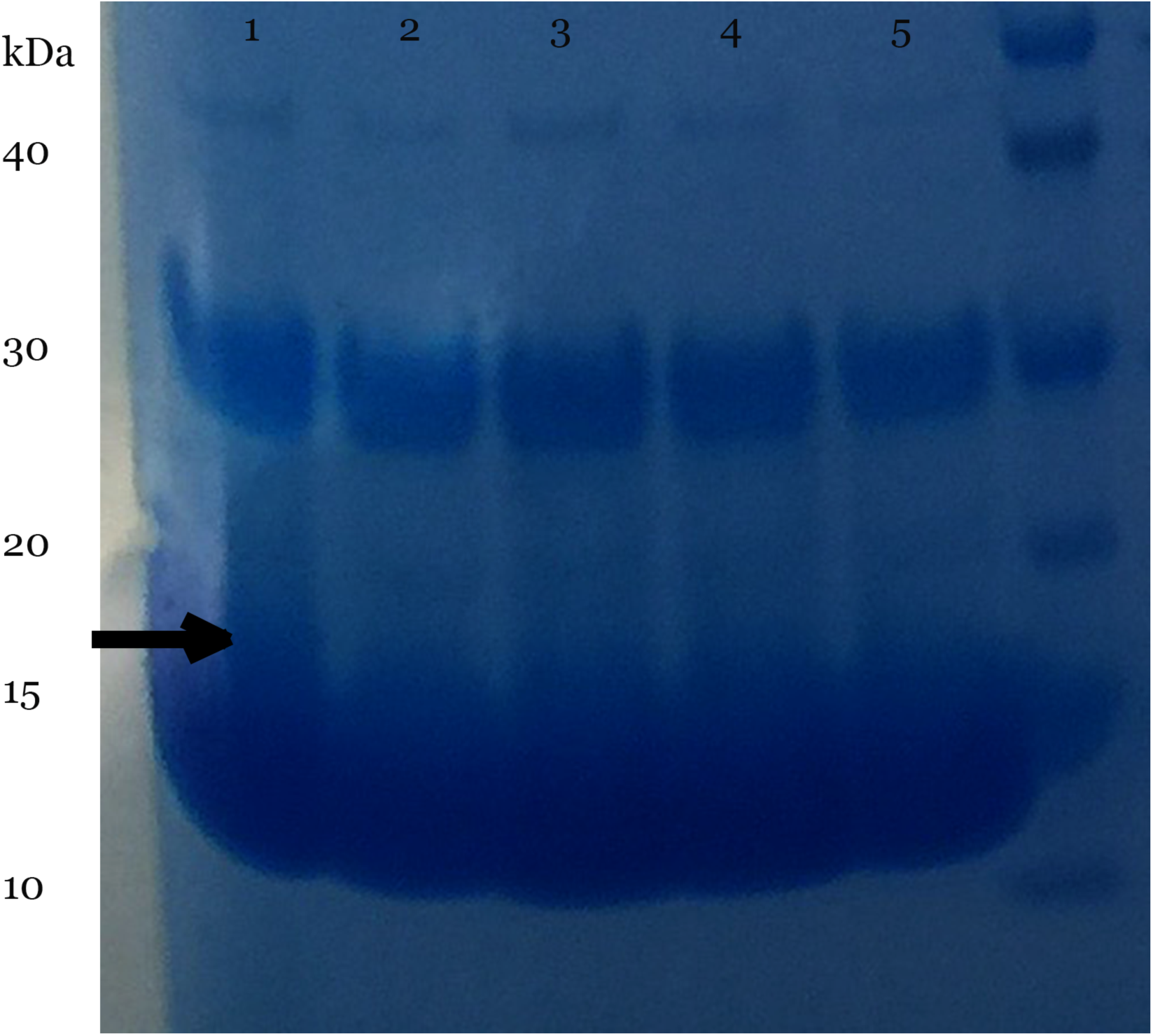
Inhibition of protein synthesis by recombinant proteins in the Rabbit Reticulate Lysate TnT^®^ Quick Coupled Transcription/Translation System. Lane 1: Control, blur at 17kDa Lane 2: Empty reaction Lane 3: PAP-S1 36% volume of reaction (50ul) (18ul of His-Tag purified PAP-S1) Lane 4: F1 36% volume (18 ul of His-Tag purified F1) Lane 5: F2 36% volume (18ul of His-Tag purified F2) ^*^All other bands are due to proteins going through the His-Tag purification column from the initial expression reaction (i.e. Globins).

The Rabbit Reticulate Lysate TnT^®^ Quick Coupled Transcription/Translation System was used to determine activity in a eukaryotic system where the same volume of PAP-S1, F1 and F2 was used (18ul of each His-Tag purified proteins). As you can see on the below figure (Figure 8), the control gave a blur at the expected size (the blur is due to globins) while all the others gave no activity whatsoever. This was expected since PAP-S1, F1 and F2 are supposed to be extremely potent in a eukaryotic system (in the order of 1nM for total inhibition). These results however are not as conclusive as the others due to their poor quality. Even after 6 washes, it was not possible to get rid of the globins that appear as a broad band migrating at 10–15kDa and, thus, interfered with the control at 17kDa but are still a good indication of the activity of the fusion proteins in a eukaryotic system.

## Conclusion

The fusion protein between Ricin A chain C terminus and PAP-S1 N terminus was observed to be functional and active in both eukaryotic and prokaryotic cell free systems with a visible increase in activity compared to the fusion protein between Ricin A chain N terminus and PAP-S1 C terminus under the same conditions. It was also observed that it was comparable in activity to PAP-S1 in a prokaryotic system and at least identical in a eukaryotic system. The expression vectors for Plant expression were thus built based on the F2 version and were found to be extremely stable. The expression vectors for the prokaryotic systems were found to be extremely unstable, and thus, a better expression vector must be developed if E. *coli* expression is sought after. There are alternative expression systems to E. *coli* available now commercially and easily accessible. Additional research should be done in order to determine both cytotoxicity and selectivity of fusion protein F2 compared to PAP-S1 as it could be a viable more potent and less cytotoxic alternative to PAP-S1 alone for both agricultural and therapeutic applications.

## Acknowledgements

N/A

